# Sex-specific effects of *in vitro* fertilization on adult metabolic phenotypes and hepatic transcriptomic and proteomic pathways in mouse

**DOI:** 10.1101/2020.09.23.309989

**Authors:** Laren Narapareddy, Eric A. Rhon-Calderon, Lisa A. Vrooman, Josue Baeza, Duy K. Nguyen, Yemin Lan, Benjamin A. Garcia, Richard M. Schultz, Marisa S. Bartolomei

**Affiliations:** Nell Hodgson Woodruff School of Nursing, Emory University, Atlanta, GA, 30322 USA; Epigenetics Institute, Department of Cell and Developmental Biology, Perelman School of Medicine, University of Pennsylvania, Philadelphia, PA, 19104 USA; Epigenetics Institute, Department of Biochemistry and Biophysics, Perelman School of Medicine, University of Pennsylvania, Philadelphia, PA, 19104 USA; Department of Biology, School of Arts and Sciences, University of Pennsylvania, Philadelphia, PA, 19104 USA

**Keywords:** Assisted Reproductive Technologies, developmental origins of health and disease, offspring, follow-up, long-term health

## Abstract

Although *in vitro* fertilization (IVF) is associated with adverse perinatal outcomes, an increasing concern is the long-term health implications. We augmented our IVF mouse model to longitudinally investigate cardiometabolic outcomes in offspring from optimal neonatal litter sizes. We found that IVF-conceived females had higher body weight and cholesterol levels compared to naturally-conceived females, whereas IVF-conceived males had higher levels of triglycerides and insulin, and increased body fat composition. Through transcriptomics and proteomics of adult liver, we identified sexually-dimorphic dysregulation of the sterol regulatory element binding protein (SREBP) pathways that are associated with the sex-specfic phenotypes. We also found that global loss of DNA methylation in placenta was linked to higher cholesterol levels in IVF-conceived females. Our findings indicate that IVF procedures have long-lasting sex-specific effects on metabolic health of offspring and lay the foundation to utilize the placenta as a predictor of long-term outcomes.

## Introduction

Over the past 40 years, assisted reproductive technologies (ART), including *in vitro* fertilization (IVF), have become a highly successful treatment for infertility. Globally, there has been a consistent and steady increase in the use of ART ^1^, now contributing to over 9 million births worldwide ^2^. Although the majority of IVF births are uncomplicated, IVF is associated with adverse perinatal outcomes, including higher risk of congenital anomalies, preterm birth, low birth weight, perinatal mortality, small for gestational age, and imprinting disorders, as well as hypertensive and placental disorders during pregnancy ^3–8^.

Beyond perinatal outcomes, it is critical to consider the success and safety of IVF within the context of developmental origins of health and disease (DOHaD). DOHaD posits that suboptimal environments and exposures during critical periods of embryo development can predispose individuals to health complications later in life ^9^. An unavoidable circumstance is that IVF procedures and manipulations occur during preimplantation development, a period of tightly coordinated physiological and epigenetic changes. IVF opens this critical window of embryo development to a synthetic, ex-vivo environment that can perturb the coordination of embryo development in ways that may affect health and disease later in life. Because IVF has only been in the clinical setting for just over 40 years, the current epidemiological evidence is limited. Nevertheless, subclinical indicators of cardiometabolic alterations have been identified in human cohort studies of children and young-adults conceived with IVF ^8,10^.

Investigating the effects of ART in fertile mouse models circumvents issues raised by underlying infertility and thereby affords the ability to dissect the effects of individual or collective ART procedures on long-term health outcomes. In agreement with human cohort studies, mouse models also demonstrate an array of cardiometabolic alterations in offspring conceived with IVF and other ART procedures. Long-lasting effects of IVF observed in mouse models, include increased body weight, increased fasting glucose, impaired glucose tolerance, insulin resistance and higher blood pressure ^11–17^, some of which occur in a sex-specific manner. However, the evidence from mouse models has yet to define clear and consistent phenotypes or contributing mechanistic pathways.

Here, we exploit our mouse IVF model with a neonatal fostering paradigm, which controls for neonatal litter size, to conduct longitudinal metabolic assessments of IVF- and naturally-conceived offspring between 12-weeks and 39-weeks-of-age. As early as 12 weeks-of-age, we detect distinct, sex-specific phenotypes that persist through adulthood. Examination of adult hepatic transcriptome and proteome identified pathways involved in these sex-specific phenotypes, in particular, sexually-dimorphic dysregulation of sterol regulatory binding element proteins (SREBPs). We also find that global dysregulation of DNA methylation in the placenta is linked to the IVF-female phenotype.

## Results

### IVF produces sexually dimorphic metabolic phenotypes

To determine the effects of IVF on metabolic phenotypes throughout the lifespan, we performed longitudinal metabolic phenotyping on IVF- and Nat-offspring. An important confounder of adult metabolic phenotypes is neonatal litter size. In particular, neonates nursed in small litters (3-4), which are commonly used in mouse models of ART, show increased body weight and adiposity in adulthood compared to neonates nursed in litters of 6-10 ^18^. We controlled for the effects of neonatal litter size on adult metabolic phenotypes by maintaining an average total litter size (native plus foster neonates) of 10 neonates throughout nursing. Then, metabolic panels were performed at 12-, 16-, 20-, 24-, 28- and 39-weeks-of-age which consisted of fasting serum measures of total cholesterol, triglycerides and insulin along with glucose tolerance testing. All fasting serum measures and statistics are displayed in Supplemental Table 1.

As early as 12-weeks we identified sex-specific differences in metabolic phenotypes. IVF-females were significantly heavier at all timepoints relative to Nat-females (Figure 1A), and the difference became more pronounced with age. IVF-females also had consistently higher cholesterol than Nat-females at all time points except for 28-weeks (Figure 1B). Nat-females demonstrated higher triglycerides and glucose area under the curve (AUC) at 16-weeks, and higher glucose at 20-weeks (Figures 1C, 1F and 1E), but these differences were transient. Although the expected age-related increase in insulin levels was observed over time in both IVF- and Nat-females, the IVF-females demonstrated significantly higher insulin at 39-weeks (Figure 1D).

**Figure 1.**
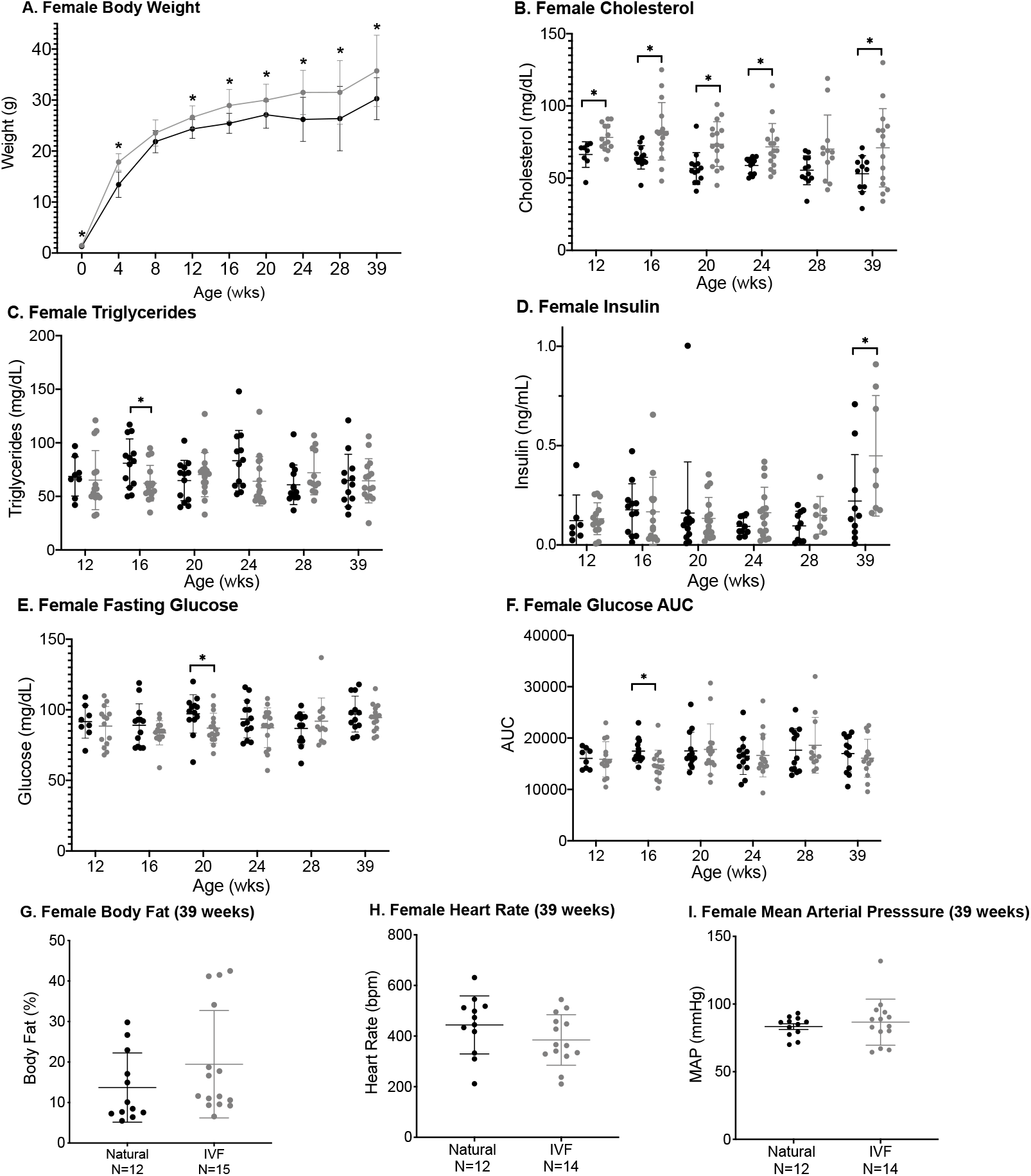
Female metabolic phenotypes in naturally-conceived and IVF-conceived offspring through 39-weeks-of-age. Data from each time point are from n=8-12 Nat-females and n=14-16 IVF-females. Body weights were measured in un-fasted mice throughout the study (**A**). At 6 timepoints, total cholesterol (**B**), total triglycerides (**C**), insulin (**D**), and glucose (**E**) were measured after 6-hours of fasting. IPGTTs were also performed at these same timepoints and analyzed by area-under-the-curve (AUC) (**F**). At the terminal timepoint (i.e. 39-weeks), body composition was measured by nuclear magnetic resonance (**G**) and heart rate (**H**) and blood pressure (**I**) were measured by non-invasive tail-cuff. Black and gray dots represent Nat-concieved and IVF-conceived females, respectively. Lines represent the mean and error bars represent the standard deviation. Statistical significance was determined using two-tailed student’s t-test. P-value <0.05 was considered significant (*).

Unlike females, we found no significant differences in body weight at the metabolic panel timepoints (Figure 2A) or in cholesterol between IVF- and Nat-males except at 39-weeks (Figure 2B). In contrast to females, IVF-males showed significantly higher triglyceride levels at 12-, 16-, 20- and 28-weeks (Figure 2C). For insulin, IVF-males displayed significantly higher levels than Nat-males at 20-, 24-, 28- and 39-weeks (Figure 2D). IVF-males also demonstrated significantly higher fasting glucose at 16- and 24-weeks, but no differences in glucose homeostasis as analyzed by AUC (Figures 2E and 2F).

**Figure 2.**
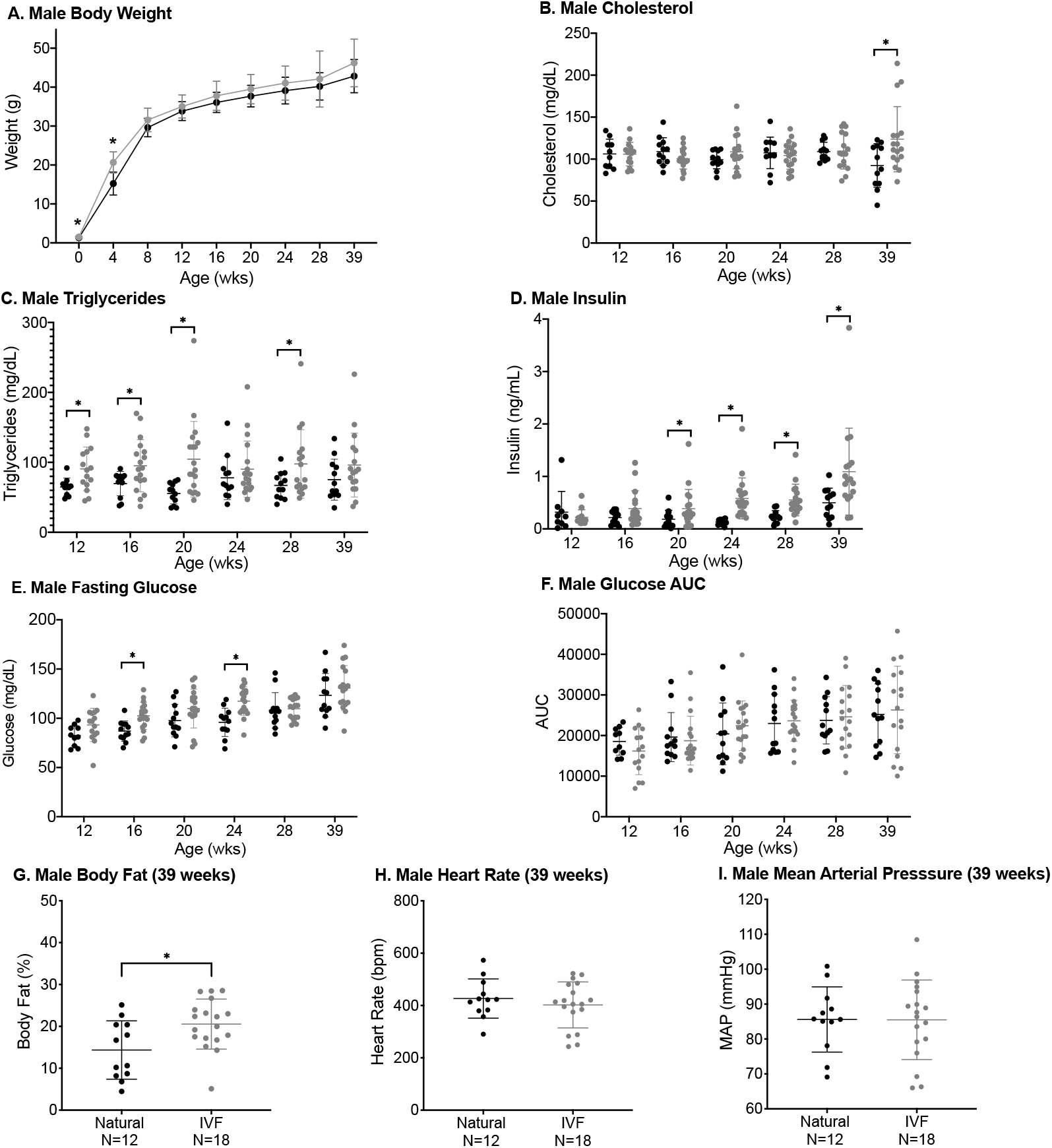
Male metabolic phenotypes in naturally-conceived and IVF-conceived offspring through 39-weeks-of-age. Data from each time point are from n=9-13 Nat-males and n=15-19 IVF-males. Body weights were measured in un-fasted mice throughout the study (**A**). At 6 timepoints, total cholesterol (**B**), total triglycerides (**C**), insulin (**D**), and glucose (**E**) were measured after 6-hours of fasting. IPGTTs were also performed at these same timepoints and analyzed by area-under-the-curve (AUC) (**F**). At the terminal timepoint (i.e. 39-weeks), body composition was measured by nuclear magnetic resonance (**G**) and heart rate (**H**) and blood pressure (**I**) were measured by non-invasive tail-cuff. Black and gray dots represent Nat-concieved and IVF-conceived males, respectively. Lines represent the mean and error bars represent the standard deviation. Statistical significance was determined using student’s t-test. P-value <0.05 was considered significant (*).

To further characterize the observed metabolic phenotypes we measured body composition, blood pressure and heart rate at 39-weeks, which was the terminal timepoint for this study (Supplemental Table 2). We found no difference in body fat percentage between IVF- and Nat-females (Figure 1G). Despite no significant difference in body weight at 39-weeks, IVF-males demonstrated significantly higher body fat percentage compared to Nat-males (Figure 2G). No differences were found in either sex for heart rate, mean arterial pressure (Figures 1H-I and 2H-I) or systolic and diastolic blood pressures.

### Sexually-dimorphic effects of IVF on metabolic phenotypes are reflected in the transcriptome and proteome of adult liver

#### Adultx liver transcriptome

To identify molecular mechanisms underlying the metabolic phenotypes, we performed RNA-seq on whole liver collected from IVF and Nat offspring at 39-weeks. Alignment of reads to reference sequence and number of mapped reads per sample are displayed in Supplemental Table 3. As the liver displays sexual-dimorphism^19,20^ we analyzed female and male liver separately. Principle Component Analysis (PCA) using the expression profile of all genes did not show separation between female IVF and Nat samples, whereas males showed separation on PC2 (Supplemental Figure 1A and 1C). Although these findings indicate that expression differences between groups are small compared to individual differences, using a cutoff FDR < 0.05 we identified 12 differentially-expressed genes (DEGs; 7 upregulated and 5 downregulated) between IVF- and Nat-females, and 17 DEGs (9 upregulated and 8 downregulated) between IVF- and Nat-males (Supplemental Figure 1B and 1D).

Based on the observed sexually-dimorphism in cholesterol and triglycerides phenotypes, we explored pathways regulated by sterol regulatory element binding proteins (SREBPs), which act as master regulators of cholesterol and fatty acid synthesis in liver ^21,22^. Of the three mammalian isoforms, *Srebp-2* preferentially activates genes involved in cholesterol metabolism, whereas *Srebp-1c* preferentially activates genes involved in fatty acid and triglyceride metabolism. Although SREBPs were not detected in our RNA-seq data, presumably reflecting very low abundance, heatmaps of genes within the *Srepb-2* regulated pathway showed higher expression in IVF-female mice (Figure 3A; unadjusted-p <0.05). No differences were observed in the *Srebp-1c* pathway (Supplemental Figure 2). Within this small sub-sample, expression differences in the *Srepb-2* pathway did not remain significant upon correcting for multiple comparisons (FDR <0.05).

**Figure 3.**
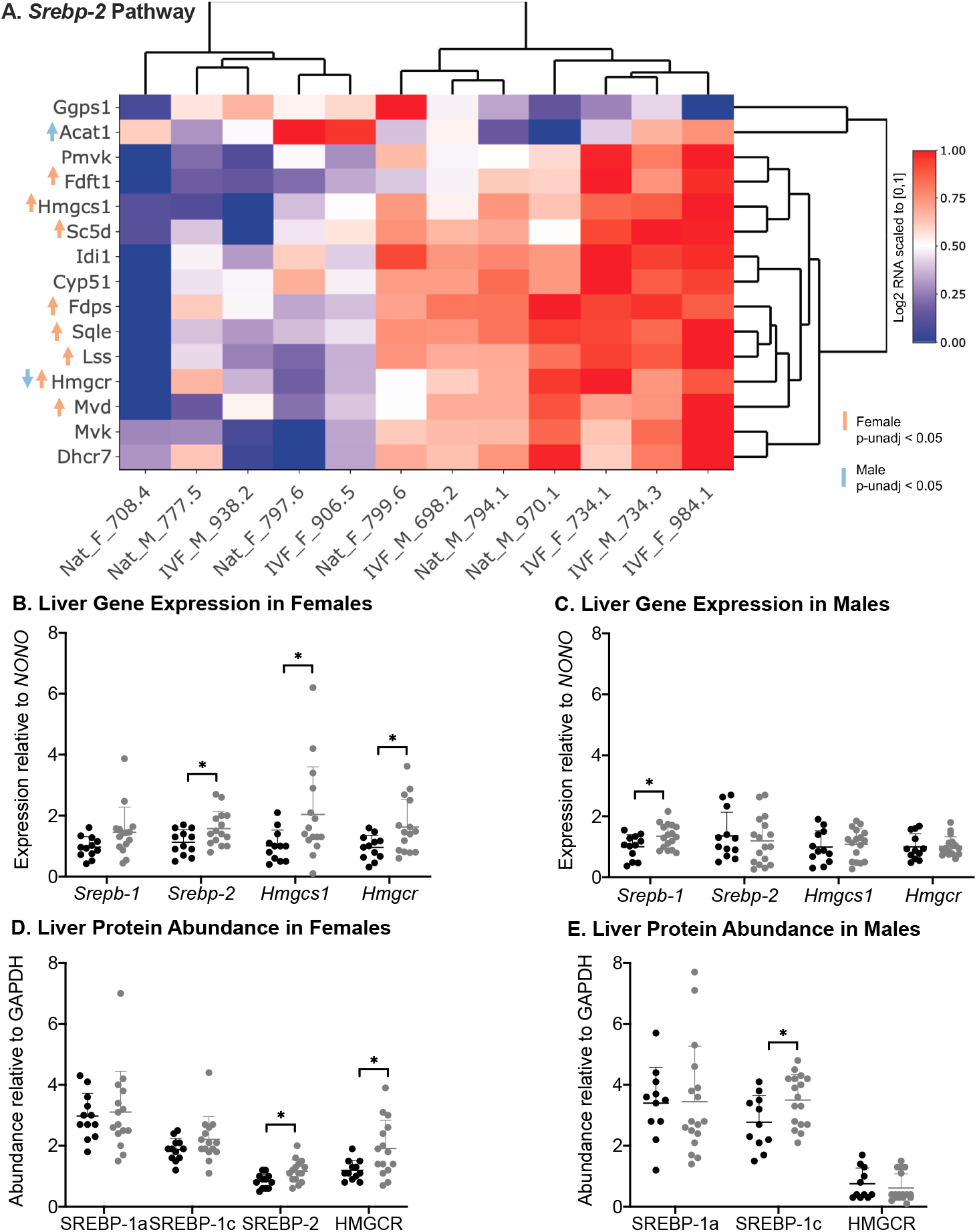
Gene expression and protein levels in 39-week liver of naturally-conceived and IVF-conceived offspring. The heatmap demonstrates log2-transformed expression levels obtained from RNA-Seq of genes within the *Srebp-2* pathway (**A**). mRNA expression levels of *Srepb1, Srebp2, Hmgcsl, Hmgcr* obtained through real-time PCR for females (**B**) and males (**C**). Downstream protein levels of SREPB isoforms 1a and 1c, SREBP-2 and HMGCR in females (**D**) and males (**E**). Black and gray dots represent Nat-concieved and IVF-conceived males, respectively. Lines represent the mean and error bars represent the standard deviation. Statistical significance was determined using student’s t-test. P-value <0.05 was considered significant (*).

We further investigated these pathways among the entire cohort by performing real-time PCR and western blot of SREBPs and their primary targets. Again, we found sex-specific differences. Consistent with their high cholesterol phenotype, IVF-females exhibited higher transcript and protein abundance of *Srebp-2* (Figure 3B and 3D). In the cholesterol biosynthetic pathway, *Srebp-2* responsive genes include HMG-CoA synthase (*Hmgcs1*) and HMG-CoA reductase (*Hmgcr*) ^21^. Transcript and protein abundance of these enzymes were elevated in IVF-females compared to Nat-females (Figure 3B and 3D), suggesting that the *Srepb-2* pathway may contribute to the high cholesterol phenotype observed in IVF-females.

In contrast, we did not observe any differences in *Srebp-2* transcript abundance in males, but rather PCR-based analyses showed upregulation of *Srebp-1* in IVF-males (Figure 3C). When we evaluated downstream protein abundance of the two SREBP-1 isoforms (i.e, SREBP-1a and SREBP-1c), we found that SREBP-1a expression, whose target genes increase both cholesterol and triglyceride biosynthesis ^22^, did not differ between IVF- and Nat-males. However, protein expression of SREBP-1c, whose downstream effects stimulate production of triglycerides and phospholipids, was significantly higher in IVF-males compared to Nat-males (Figure 3E). These findings suggest that SREBPs are involved in development of IVF phenotypes and that the effects of IVF on the SREBP pathways are sex-specific and underlie the sex-specific phenotypes.

#### Adult liver proteome

We also used liquid chromatography mass spectrometry-based proteomics to identify protein abundance in liver collected at 39-weeks from a subset of IVF and Nat offspring (n=5/sex/group). We identified a total of 24,037 peptides with abundance above zero and a median peptide coefficient of variation (CV) of 18.22. Applying a cut-off CV of 50, the 24,037 peptides corresponded to 2,924 proteins. The heatmap in Figure 4A demonstrates significant proteins in the liver proteome identified by 2-way ANOVA.

**Figure 4.**
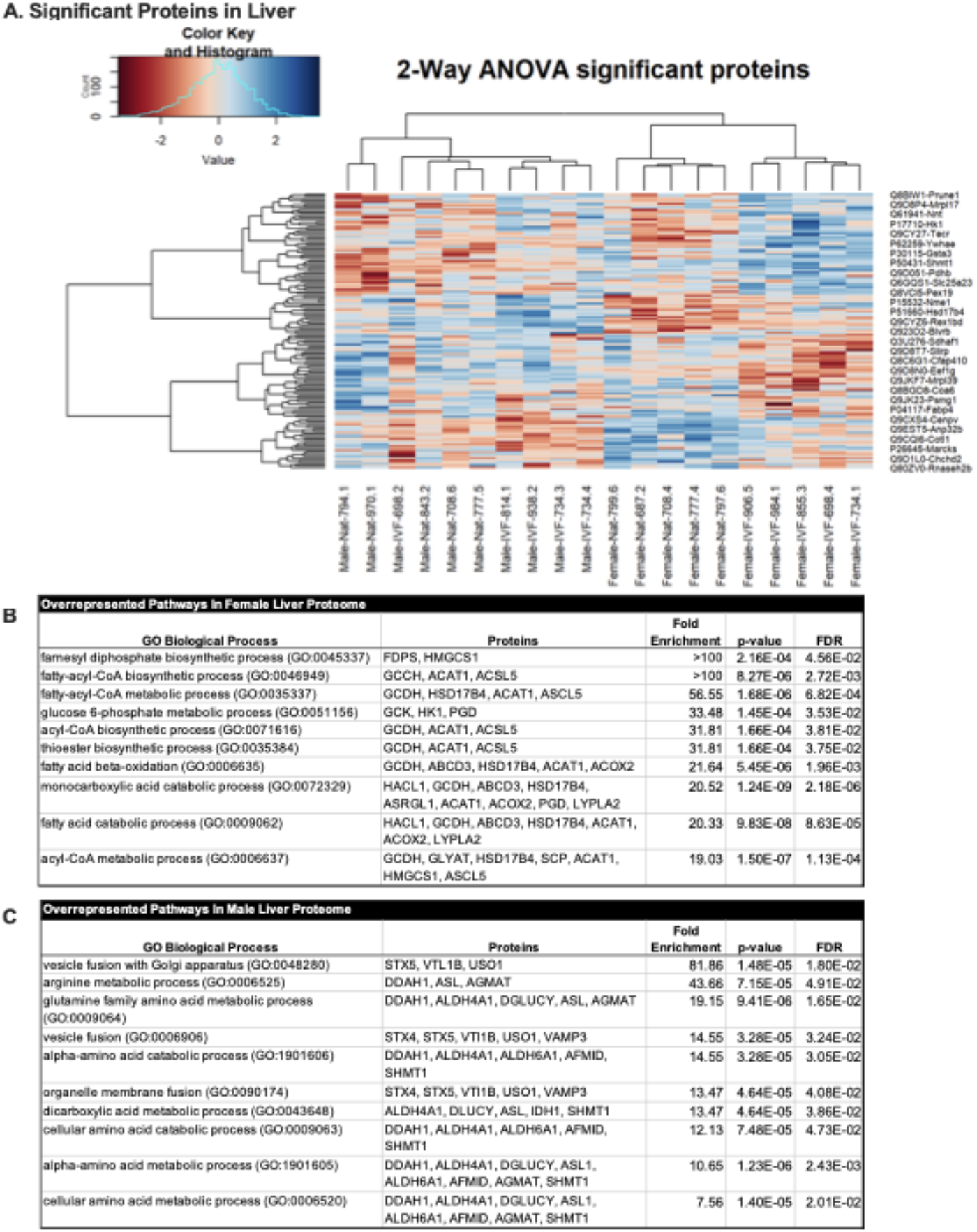
Liver proteome in 39-week naturally-conceived and IVF-conceived offspring. The heatmap demonstrates results from 2-way ANOVA of log2-transformed protein expression levels obtained from liquid chromatography mass spectrometry **(A)**. The top 100 proteins were used for overrepresentation tests in PANTHER to identify overrepresented biological processes in female **(B)** and male **(C)** liver proteome.

Two-way ANOVA identified 43 proteins with differential abundance between IVF-female and Nat-female (FDR <0.05, 20 upregulated and 23 downregulated). Using the top 100 proteins based on p-values, we performed gene ontology analysis with the PANTHER overrepresentation test. The top 10 overrepresented GO pathways in females (Figure 4B) suggest that IVF-females and Nat-females differ in pathways important for cholesterol biosynthesis (FDPS, HMGCS1), fatty acid metabolism (GCCH, ACAT1, ASCL5, GCDH, ABCD3, HSD17B4, etc.) and energy production pathways involving glucose 6-phosphate (GCK, HK1, PGD). Of particular relevance to the IVF-female phenotype of high cholesterol is the upregulation of FPDS and HMGCS1 that are involved in the farnesyl diphosphate biosynthetic process (GO:0045337), an important intermediate in conversion of mevalonate to cholesterol. Although FPDS was in the top 100 proteins identified by two-way ANOVA in females, differential abundance of FPDS did not remain significant after correction for multiple testing (FDR=0.1187). However, consistent with our evaluation of gene expression (Figure 3) and further supporting involvement of genes responsive to *Srebp-2,* HMGCS1 displayed significantly greater abundance in IVF-females compared to Nat-females (log2 fold change = 1.4062, FDR=0.0130).

In males, we identified 24 proteins with differential abundance between IVF-male and Nat-male (FDR <0.05, 11 upregulated and 13 downregulated). Again, we performed an overrepresentation test in PANTHER using the top 100 proteins; Figure 4C shows the top 10 overrepresented GO pathways. The identified pathways suggest potential differences in vesicle fusion processes (STX5, VTI1B, USO1) and amino acid metabolism (DDAH1, ALDH4A1, DGLUCY, ASL, AGMAT, SHMT1, etc.) between IVF- and Nat-males. For proteins involved in amino acid metabolism, several remained significantly different after correction for multiple testing (STX5, PDXDC1, IDH1, DGLUCY, and CYP2C55; FDR < 0.05), indicating that hepatic amino acid metabolism is one of the most pronounced differences between IVF- and Nat-males. As excess amino acids can be interconverted and metabolized directly into precursors for production of glucose and fatty acids, our findings suggest that disruption of amino acid metabolism in IVF-males may be a potential contributor to the IVF-male phenotype characterized by high triglycerides.

#### Adult liver histology

Given the sexually-dimorphic differences observed in liver, we performed liver histology in a random subset of offspring to assess hepatocellular swelling, clearing, atrophy, vacuolation, karyomegaly and necrosis as well as sinusoidal inflammatory cell aggregates. Liver histology assessments performed at 39-weeks did not reveal marked differences. Many of the lesions noted in the liver samples are considered common background findings, such as the extramedullary hematopoiesis, portal inflammatory cell infiltrates and inflammatory cell aggregates within the sinusoids with or without individual hepatocellular necrosis. These traits did not differ between sexes or groups. Both groups and sexes also exhibited expected age-related changes with no differences regardless of individual phenotypes.

### Replication of findings in 12-week cohort

To replicate our findings in an independent cohort as well as determine if differences in liver proteins could be detected early on, we generated an additional cohort of IVF- and Nat-offspring and assessed them at 12 weeks-of-age. Similar to our findings from the original cohort, IVF-females from the 12-week-cohort had significantly higher total cholesterol (p=0.0022; Supplemental Figure 3A). For this cohort, we quantified HDL and combined LDL/VLDL fractions and found that both fractions were higher in IVF-females compared to Nat-females (HDL: p=0.0174; LDL/VLDL: p=0.0418). As with the original cohort, the 12-week cohort did not exhibit a significant difference in triglycerides between IVF- and Nat-Females (p=0.682; Supplemental Figure 1B). Like our original cohort, IVF-males showed no differences in total, HDL and LDL/VLDL cholesterol (Supplemental Figure 3C), but elevated triglycerides were again evident at 12-weeks (p=0.0155; Supplemental Figure 3D).

We also confirmed sexually-dimorphic protein expression of SREBPs and HMGCR. Combined with our results from 39-week liver, elevation of SREBP-2 and HMGCR in 12-week IVF-females (Supplemental Figure 3C) indicates that this pathway of cholesterol biosynthesis is disrupted early on and disruption persists through adulthood. Unlike our at 39-week cohort, SREBP-1 was elevated in 12-week IVF-females compared to Nat-females. As SREBP-1a and 1c were not quantified separately, it is unclear which isoform contributes to this increase. SREBP-1c preferentially activates fatty acid and triglyceride synthesis, SREBP-1a activates both triglyceride and cholesterol synthesis. Although we detected no differences in either isoform in 39-week liver, based on the high cholesterol phenotype, it is possible that the elevation may be transient and due to SREBP-1a.

As expected, no differences were observed in SREBP-2 or HMGCR in liver of 12-week males (Supplemental Figure 3F). SREBP-1 was significantly higher in IVF-males compared to Nat-males and given the results from our 39-week cohort, we hypothesize that this increase is driven by the preferential activator of triglyceride biosynthesis, SREBP-1c.

### IVF produces sexually dimorphic placental epigenotypes

In previous work, we and others have shown that IVF procedures induce placental epigenotypes characterized by reduced global DNA methylation and loss of monoallelic expression at imprinted genes ^23–29^. To obtain placental tissues and deliver neonates at similar developmental stages, Nat offspring were delivered at E18.5 and IVF offspring at E19. To assess whether placental epigenetic changes were sex-specific, we used LUMA (see methods)^23,30,31^ to measure global DNA methylation and pyrosequencing to measure DNA methylation at the regulatory imprinting control regions (ICRs) of four imprinted genes that are involved in regulation of placental and fetal development. As the ICRs of imprinted genes exhibit methylation on only one allele, non-allele-specific measurement yields roughly 50% methylation.

We performed non-allele-specific measurement of DNA methylation at the ICRs of two paternally methylated ICRs--*H19/Igf2* and *IgDMR* (the ICR for the *Dlk1/Gtl2* imprinted cluster)—as well as two maternally methylated ICRs—*KvDMR1 (the* ICR for the *Kcnq1* cluster) and *Peg3.* We found sex-specific differences in global and ICR DNA methylation between IVF and Nat (Supplemental Table 4). IVF-females demonstrated significantly lower DNA methylation globally and at the ICRs of *H19/Igf2* and *Peg3,* but no differences at *KvDMR1* and *IgDMR* compared to Nat-females (Figure 5A). Conversely, IVF-males had lower DNA methylation at the *Peg3* ICR but no differences globally or at the *H19/Igf2* ICR, *KvDMR1,* or *IgDMR* compared to Nat-males (Figure 5B). These results support previous findings of loss of imprinting in placentas of IVF-conceived pregnancies and provides new evidence that these methylation changes are sex- and gene-specific.

**Figure 5.**
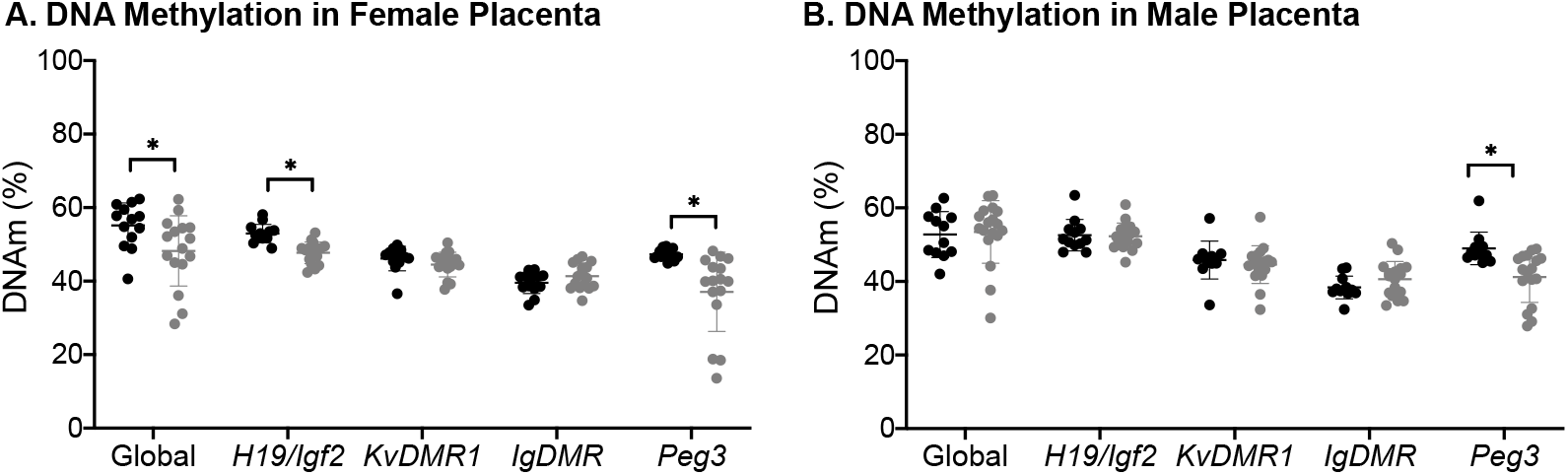
DNA methylation in naturally-conceived and IVF-conceived placentas. Each data point represents an individual female placenta from n=13 Nat-females and n=16 IVF-females. Luminometric methylation assay was used to measure global DNA methylation. Bisulfite pyrosequencing was used to measure Imprinting control region DNA methylation in placenta at *H19/Igf2, KvDMR1, IgDMR,* and *Peg3* in female (**A**) and (**B**) male placenta. Black and gray dots represent Nat-concieved and IVF-conceived males, respectively. Lines represent the mean and error bars represent the standard deviation. Statistical significance was determined using student’s t-test. P-value <0.05 was considered significant (*).

### Average cholesterol of IVF females is correlated with global DNAm

Placental epigenetic marks regulate fetal development and variations in these marks have been proposed as biomarkers of early life growth and long-term health outcomes. To evaluate the potential of placental epigenetics marks as predictive biomarkers, we assessed whether the identified placental epigenotypes were associated with the sex-specific offspring outcomes. Accordingly, we used a Cesarean delivery and fostering protocol that enabled us to distinguish which placenta belonged to which offspring by permanently marking each neonate with a unique identifier at birth. We then performed correlation analysis to evaluate whether or not placental DNA methylation was associated with the long-term phenotypes observed in this study. For serum measures, we used the average of the measures over time. Upon correlation analyses (Supplemental Figure 4), IVF-females demonstrated a correlation between average cholesterol and global placental DNA methylation. No other correlations were found between other measures of placental epigenotype and offspring metabolic phenotypes.

## Discussion

Throughout the field of developmental programming, evidence showing that prenatal exposures affect males and females differently ^32,33^ continues to grow. In mouse, more than 600 transcripts are differentially expressed between the sexes as early as the blastocyst stage ^34^, suggesting that the early embryo can respond to ART procedures in a sex-specific manner. Here we show that ART procedures during preimplantation development result in differential effects based on sex and that these effects have long-lasting consequences on metabolic parameters and pathways in adulthood.

We and others find that IVF-females are heavier in adulthood compared to controls. Although we detect higher body weight at 12-weeks, Feuer et al. ^14^ did not observe higher body weight until 17-weeks. Feuer et al., also noted IVF-females exhibit increased body fat and altered glucose homeostasis, whereas we do not. Although Feuer et al., note no differences in outcomes based on litter size, differences between their findings and ours may be due to a variety of other reasons including comparison groups. Feuer et al., use a flushed blastocyst control group with litter sizes ranging from 3-8. We and others have already identified how individual ART procedures, including those used to generate a flushed blastocyst control group, introduce iatrogenic changes to fetal and placental development ^23,24,26,31^ and that the full IVF protocol demonstrates the most severe phenotypes. Therefore, our intent here was to take a clinically relevant approach by examining the full IVF protocol in comparison to natural conception.

In contrast to females, we find a novel and more severe metabolic phenotype in IVF-males characterized by high insulin, triglycerides, and body fat percentage. Abnormality in all three of these metabolic parameters occurs in development of severe metabolic disease ^35,36^. Hence, these findings are potentially clinically relevant and suggest that males conceived through IVF may be at increased risk for metabolic dysfunction. Others have found that male mice conceived through various ART procedures exhibit reduced lifespan with high fat diet ^37^, insulin resistance ^15^, altered glucose homeostasis ^15,17^, increased blood pressure ^11,12,37^, or no differences compared to controls ^13,14^.

One challenging aspect of long term outcome studies, is that litter size can contribute to differences in outcomes between studies. We maintained an average neonatal litter size of 10, because it is a standard control size in studies evaluating the impact of neonatal litter size on metabolic pheontypes ^18^. Litter sizes reported in the studies cited here range from 2-8, noting that litter sizes <5 are considered small. Small neonatal litter size is associated with offspring that exhibit increased food intake during lactation, leading to increased body weight and adiposity in adulthood ^18^. Thus, similar to dietary challenge models, studies of ART that use small litter sizes could present a post-natal “challenge” of increased food availability and intake that is inherent to the study design and potentially confounds the effects of ART. Other factors contributing to variations in outcomes between studies include differences in genetic background, comparision groups, IVF protocols, fasting protocols, and the use of high-fat dietary challenges.

To our knowledge, this is the first study in an animal model of ART to longitudinally evaluate triglyceride and cholesterol levels in IVF-conceived offspring and moreover, to assess sex differences, which we find. The few human studies evaluating triglyceride and cholesterol in children and young adults conceived with IVF have not demonstrated adverse lipid profiles ^17,38^. In fact, one study of children between the ages of 4- and 10-years-old demonstrated a favorable profile of lower triglycerides and increased HDL cholesterol ^39^. Many of the human studies assessing metabolic outcomes in IVF-conceived children are limited by lack of longitudinal evaluation and small sample sizes that do not allow for sex-specific analyses, thus, masking any sex-specific effects that may be present. Whereas studies in young adults and children conceived with IVF are essential, an important finding from mouse models is that some phenotypes resulting from IVF may not present until early adulthood or later. Taken together, our findings highlight the necessity of longitudinal assessment across several parameters of cardiometabolic health including lipid profiles, and the need to evaluate these parameters in a sex-specific manner throughout life.

Because we found novel sex-specific differences in cholesterol and triglyceride levels, we used transcriptomics and proteomics to compare how IVF affects molecular mechanisms in the liver between females and males. We explored pathways regulated by sterol regulatory element binding proteins (SREBPs), transcription factors that act as master regulators of cholesterol and fatty acid synthesis in liver. Of the three mammalian isoforms, SREBP-2 preferentially activates genes involved in cholesterol metabolism, whereas SREBP-1c preferentially activates genes involved in fatty acid and triglyceride metabolism. Given these differences, we hypothesized that we would see differences in these pathways based on sex.

Consistent with the high cholesterol phenotype found in IVF-females, transcriptomic analysis of the SREBP-2 pathway shows several genes that trended toward upregulation in females conceived through IVF (*Fdft1, Hmgcs1, Sc5d, Fdps, Sqle, Lss, Hmgcr,* and *Mvd*). Likewise, pathway analysis of the liver proteome from a subset of individuals indicates overrepresentation of proteins involved in cholesterol biosynthesis (GO:0045337; FDPS and HMGCS1); we confirmed upregulation of SREBP-2 and its main targets (i.e., HMGCS1 and HMGCR) in the entire cohort. Combined, these findings suggest that IVF is capable of inducing long-lasting effects on cholesterol homeostasis through upregulation of SREBP-2 and that these effects are specific to females.

Beyond the SREBP pathways, our analyses of the female liver proteome also identified alterations in pathways of fatty acid metabolism (GO:0046949, GO:0035337, GO:0071616, GO:0006635, GO:0009062, GO:0006637). Although phenotypes differ, liver proteomics assessing *ob/ob* and *db/db* mice ^40^, high-fat diet in rat ^41^, and cholesterol-induced atherosclerosis in rabbit ^42^ also show disturbances in lipid metabolism. With the diverse role of fatty acid metabolism, including dependence on substrates that are also involved in cholesterol biosynthesis, it is not surprising that fatty acid metabolism is commonly affected in various models of metabolic dysfunction, including IVF.

In line with the high triglyceride levels exhibited in IVF-males, we find dysregulation of hepatic SREBP-1c. Although our exploratory transcriptomic assessment of the SREBP-1c pathway did not show differences in gene expression between naturally- and IVF-conceived males, locus-specific assessment of SREBP-1c in the entire cohort showed increased abundance of SREBP-1c. SREBP-1c acts as the master regulator of lipogenesis by preferentially activating the entire complement of genes involved in fatty acid and triglyceride synthesis ^22,43,44^. Insulin activates SREBP-1c by increasing transcription ^45^ and by increasing the conversion of the inactive precursor to the active nuclear form^46^. Like the IVF-males of this study, mouse models of insulin resistance, such as high fat diet ^47^ and *ob/ob* mice ^48^ have also demonstrated higher levels of SREBP-1c. Our findings suggest that the higher insulin levels associated with IVF may trigger signaling events that increase hepatic SREBP-1c and lipogenesis, ultimately perpetuating the high serum triglycerides in IVF-males. IVF-males demonstrate higher triglyceride levels as early as 12-weeks and higher insulin levels are not detected until 20-weeks. Even so, it is possible that the insulin-dependent SREBP-1c pathway may be more sensitive to insulin in IVF-males even in the absence of higher insulin levels at the earlier timepoints.

Although the majority of offspring conceived with IVF are generally healthy, our work and that of others suggest that those who develop adverse metabolic outcomes may not do so until later life. As such, an important task in optimizing IVF outcomes is to identify biomarkers of long-term health. The placenta has been proposed as an easily accessible tissue that may serve as one such biomarker at birth. Additionally, we have previously shown locus-specific effects of IVF on DNA methylation in the placenta ^23,24,31^. Here, our placental collection and fostering protocol provided the unique opportunity to assess whether placental epigenetic changes at birth could serve as biomarkers of future metabolic outcomes in offspring conceived with IVF. In correlating placental DNA methylation with sex-specific phenotypes, we found that global placental DNA methylation is associated with serum total cholesterol in females conceived through IVF. To our knowledge, this is the first study to demonstrate a link between placental DNA methylation and long-term outcomes in offspring. This finding remains to be validated in a larger cohort and in a manner that accounts for cellular heterogeneity, but provides promise for the use of placental epigenetic biomarkers to identify individuals at birth who may be at risk for adverse metabolic outcomes later at life.

Overall, the findings presented here support the growing body of evidence that IVF contributes to altered metabolic health outcomes later in life ^8,11–17^ and that this effect occurs in a sex-specific manner. Although mouse is an excellent and efficient model of investigation that allows for the IVF procedures to be isolated from underlying fertility, the findings here may be limited in their translation to human outcomes. Because oxygen, temperature, pH, humidity, light and culture conditions are not consistent across all models of IVF—nor are they standardized in the clinical setting— phenotypes and epigenotypes may be specific to these conditions, emphasizing the importance of standardization in clinical and pre-clinical settings. Moreover, our findings highlight the necessity of longitudinal and sex-specific monitoring of outcomes in individuals conceived with IVF and the need for continual efforts in optimizing IVF procedures to ensure the future health of offspring.

## Materials and methods

### Animals

CF1 female mice (Envigo), B6SJL male mice (Jackson Laboratory), and CD1 vasectomized male mice (Charles River) were housed in polysulfone cages in a pathogen-free facility on a 12-12 light-dark cycle. All animals had access to *ad libitum* water and standard chow (Laboratory Autoclavable Rodent Diet 5010, LabDiet, St. Louis, MO, USA), unless fasting for metabolic assays (discussed below). All animal work in this study was conducted with the approval of the Institutional Animal Care and Use Committee at the University of Pennsylvania (Protocol Number: 803545).

### Generation of Natural and IVF offspring

For the natural control group (Nat), naturally-cycling CF1 females were mated overnight with B6SJL males and embryonic day (E) 0.5 was noted as the day on which a copulatory plug was observed. Development occurred *in vivo* until E18.5.

Optimized protocols recommended by Jackson Laboratory^49^ were used to generate the IVF group. CF1 female mice were superovulated with 5 IU of eCG followed by 5 IU of hCG administered 48 h later. Approximately 14 h after injection of hCG, cumulus-egg complexes were collected from oviducts and IVF was conducted using EmbryoMax® Human Tubal Fluid (1x) medium (HTF, EMD MilliporeSigma, Burlington, MA, USA) with capacitated mature spermatozoa collected from the cauda epididymis and vas deferens of B6SJL male mice. After 4 h, eggs were washed in HTF media followed by culture in EmryoMax® KSOM media (1x) containing ½ Amino Acids (KSOM+AA; EMD MilliporeSigma, Burlington, MA, USA) in an incubator at optimized conditions (37°C, 5% CO_2_, 5% O_2_, 90% N_2_) under mineral oil suitabile for embryo culture (MilliporeSigma, Burlington, MA, USA). After four days of development in culture, fully-expanded blastocysts of similar morphology were briefly washed in HEPES-buffered minimum essential medium (MEM) prior to transferring to 2.5-day pseudo-pregnant recipient females using non-surgical embryo transfer (NSET) devices (Paratechs, Lexington, KY, USA). Each pseudo-pregnant female (generated by mating CF1 females with vasectomized CD1 males) received 10 blastocysts; The day of embryo transfer was defined as E4 and IVF blastocysts developed *in vivo* from E4 to E19.

### Fostering of Natural and IVF offspring

To account for differences in developmental rates between Nat and IVF fetuses, Caesaren sections were performed at E18.5 (Nat) and E19 (IVF) to obtain both the fetus and placenta. We also employed the following fostering protocol to minimize the effects of litter size on growth and adult metabolism ^18^. We generated the foster mother and her native litter by mating a CF1 female with a CF1 stud male. The foster mother gave birth to her native litter < 36 h prior to fostering IVF or Nat neonates into the native litter. To begin fostering, the foster mother was removed from the cage shortly before Caesarean delivery of the IVF and Nat neonates, which were quickly weighed and given a unique tattoo identifier to match neonates with their placentas. The neonates were then immediately integrated into the foster mother’s native litter. The native litter was culled so that the average total litter size (native plus foster neonates) was 10 neonates/foster mother. The foster mother was returned to the cage as soon as the fostering procedure was complete.

We produced a total of 13 IVF-females and 19 IVF-males from 19 IVF-litters and 13 Nat-females and 12 Nat-males from 9 Nat-litters. A replication cohort was generated in the same manner and aged to 12-weeks. At 12-weeks, metabolic phenotyping was performed on 16 IVF-females and 12 IVF-males from 7 litters and 12 Nat-females and 12 Nat-males from 4 litters.

### Metabolic Phenotyping

#### Body weight

Using a calibrated digital scale, body weights were measured at birth and every week from 1-week through 39 weeks-of-age.

#### Intraperitoneal glucose tolerance test (IPGTT)

Beginning at 12 weeks-of-age, IPGTTs were performed by an expert technician blinded to the experimental group of each mouse every 4 weeks until and 28-weeks of age, and again at 39-weeks (i.e., end of life). In line with optimal conditions for assessing glucose tolerance^50^, mice were fasted for 6 h during the light cycle (0800-1300) on alpha-dri bedding with *ad libitum* access to water. IPGTTs were performed by an expert technician blinded to the experimental group of each mouse. Baseline (*t*=0-min) blood glucose levels were obtained by tail snip from conscious unrestrained mice using a hand-held glucometer (*ReliOn*). Subsequently, the mice received an IP injection of 20% dextrose (2 g/Kg body weight) and blood glucose levels were measured at *t*=15-, 30-, 60- and 120-minutes post-injection.

#### Serum hormone and lipid assay analysis

During the IPGTT at *t*=0 min, whole blood was collected from the tail snip in 6-h fasted mice, placed into appropriate serum separator tubes, centrifuged and serum aliquoted and stored at −20°C for subsequent assay measurements. All assays were performed by an expert technician from the Radioimmunoassay and Biomarker Core (UPenn) who was blinded to the experimental group of each mouse. Insulin analysis was performed using a mouse insulin ELISA assay from ALPCO (Salem, NH, USA). Lipids were assayed from serum with enzymatic colorimetric assays using the following reagent kits: Triglycerides and total cholesterol from Stanbio (Boerne, TX, USA). For the 12-week cohort, total, HDL, and LDL/VLDL were obtained using the Abcam HDL and LDL/VLDL Cholesterol Assay Kit (ab65390; Cambridge, UK).

#### Body composition, blood pressure, and heart rate measurements

At 39-weeks (i.e., end of life), body weight and body composition were measured in ad-lib fed conscious mice using EchoMRI™ 3-in-1 system nuclear magnetic resonance spectrometer (Echo Medical Systems, Houston, TX, USA) to determine whole body lean and fat mass. Subsequently, systolic and diastolic blood pressure and heart rate were measured by non-invasive tail-cuff volume pressure recording (VPR) using the CODA® 8 BP System (Kent Scientific Corporation, Torrington, CT, USA). Briefly, mice were placed into an animal holder on a warming platform and tails were threaded into an O-cuff and then a VPR cuff. Before initiating the blood pressure measurement, a non-contact infrared thermometer was used to confirm that the temperature at the base of the tail was 32-35°C. Subsequently, two runs of 30 measurement cycles were performed to obtain measurement averages. All assays were performed by an expert technician blinded to the experimental group of each mouse.

### Tissue collection

At the time of fostering (see above), placentas were dissected, cleaned and weighed, and then bisected through the umbilical attachment. Half of the placenta was snap frozen in liquid nitrogen and stored at −80°C until processed for DNA and RNA isolation (see below). The other half was fixed in 10% phosphate-buffered formalin for future histological analyses.

At 12-weeks and 39-weeks, mice were euthanized via CO2 and cervical dislocation. All vital organs, including liver, were collected and snap frozen in liquid nitrogen. A portion of the left lobe of the liver was fixed in 10% phosphate-buffered formalin for histological analyses.

### DNA and RNA isolation

DNA and RNA were simultaneously isolated from one-quarter of snap frozen placentas and snap frozen liver by using phenol-chloroform extraction and TRIzol (Invitrogen, CA, USA), respectively, as previously described^23^.

### Bisulfite Pyrosequencing and Luminometric Methylation Assay (LUMA)

Using 1 μg of bisulfite-treated DNA, DNA methylation was measured at the imprinting control region (ICR) of several imprinted genes by bisulfite pyrosequencing as previously described ^23,30^.

Global DNA methylation at repetitive elements was measured by LUMA using 500 ng of genomic DNA as previously described ^23,30^.

### RNA sequencing

We performed RNA sequencing on a random subset of liver samples from the 39-week cohort (n=3/sex/group, from 11 litters). Total RNA (4 μg) was used as input for poly-A selection and mRNA-sequencing library synthesis using KAPA mRNA-Seq library synthesis kit and KAPA Single-Indexed adapter kit (Kapa Biosystems, Wilmington, MA, USA). Library quality control was conducted using Agilent 2100 Bioanalyzer system, High Sensitivity dsDNA Qubit system and NEBNext Library Quantification Kit (New England Biolab, Ipswich, MA, USA). Sequencing was performed on Nextseq 500 platform (Illumina, San Diego, CA).

RNA-seq reads were trimmed for quality using Trimmomatic (version 0.32) ^51^. Illumina TruSeq3-PE primers, leading and trailing low quality (below quality 3) and N base calls were trimmed, and reads were scanned using a 4-bp sliding window and trimmed when the average quality per base dropped below 15. Reads with at least 30-bp in length following this quality control process were retained and used for subsequent analyses. The resulting reads were then aligned to mm10 reference genome assembly using STAR (version 2.3.0e) ^52^, allowing read pairs to align no farther than 2000-bp apart and keeping alignments with a mapping quality score MAPQ greater than 1 for genome-wide transcript counting. FeatureCounts (version 1.5.0) ^53^ was used to generate a matrix of mapped fragments per RefSeq annotated gene. Numbers for total sequenced, aligned and counted reads for each sample are listed in Supplemental Table 3. Analysis for differential gene expression was performed using DESeq2 ^54^ with the cutoff of FDR <0.05 (Supplemental Dataset 1). Clustering and heatmap of differentially expressed genes were generated using heatmaply R package. The raw sequencing data reported in this work have been deposited in the NCBI Gene Expression Omnibus under accession number GSE158029 (*not yet published*).

### Quantitative Mass Spectrometry

#### Sample Preparation

The same subset of samples used for RNA sequencing was also used for liver proteomics with the addition of 2 random samples/sex/group (n=5/sex/group). A section of liver tissue was lysed by adding 2X packed tissue volumes of lysis buffer (6 M guanidine-HCl, 100 mM TEAB pH=8.0) followed by sonication using a probe sonicator at 4°C. Protein estimation was performed using a Bradford assay (Bio-Rad) with BSA as the standard. A 50-μg aliquot of each sample was volume-adjusted to 25 μL (with 6 M guanidine-HCl, 100 mM TEAB pH=8.0, 10 mM DTT), denatured and reduced while mixing on a Thermomixer C at 60°C for 20 min followed by cysteine alkylation using 40 mM iodoacetamide in the dark for 20 min. Prior to proteolytic digestion, guanidine-HCl wad diluted to 1 M using 100 mM TEAB pH=8.0. Samples were digested overnight at ambient temperature using 1 ug of Trypsin (Promega, Madison, WI, USA). Prior to LCMS, digested peptides were desalted using in-house made desalting-tips ^55^, dried using vacuum centrifugation and resuspended into 50 μL of 2% acetonitrile, 0.1% formic acid.

#### Liquid Chromatography

Approximately 1 μg of peptides were loaded and pre-concentrated onto an Acclaim™ PepMap™ 100 C18 pre-column (0.3 mm x 5 mm, 5um) using 0.05% trifluoroacetic acid in water followed by loading onto an analytical, in-house-packed fused silica capillary, C18 column (75 μm x 30 cm, 2.4 μm Reprosil-Pur Dr Maisch GmbH) using a Dionex Ultimate 3000 RSLCnano high-performance liquid chromatographic system at a flow rate of 300 nL/min. Mobile phase A consisted of an aqueous solution of 0.1% formic acid and mobile phase B as 0.1% formic acid in 80% acetonitrile. Peptides were separated using a linear gradient from 5% to 35% B over 90 min followed by a gradient ramp to 60% B over 10 min. The system was washed in 95% B for 15 min and re-equilibrated to initial conditions for 19 min.

#### Mass Spectrometry

Eluting peptides were injected into a Thermo Q-Exactive HFX and acquired using data-independent acquisition (DIA) with the chromatogram library workflow as previously described ^56,57^. An MS survey scan was collected in centroid mode for the mass range of 385 m/z to 1015 m/z (60,000 MS1 resolution, automatic gain control (AGC) 1E6 ions, 60 ms max ion injection time). The MS1 scan was followed by precursor isolation windows for fragmentation with the normalized collision energy (NCE) set to 27 and the default charge state was set to 3. Proteome profiling was performed using single-injection DIA mass spectrometry (15,000 fragment ion resolution, 1E6 AGC, 20 ms max IT) using 75-8 m/z staggered precursor isolation windows (4 m/z after demultiplexing) with optimized window placement ^58^. The precursor isolation windows were generated using EncyclopeDIA’s built-in window scheme wizard ^57^ using the following settings Window scheme – staggered/overlapping DIA; Phospho enriched – FALSE; number of windows – 75; start m/z – 400; stop m/z – 1000; margin width – 0.

To generate the chromatogram library, an aliquot of all peptide samples was pooled and analyzed using six gas phase fractionation DIA (GPF-DIA) fractions. Each of the six GPF-DIA injections were acquired using different precursor mass ranges (500-600 mz, 600-700 m/z, 700-800 m/z, 800-900 m/z 900-1000 m/z) at 30,000 MS2 resolution using 25-4 m/z staggered precursor isolation windows (2 m/z after demultiplexing) with optimized window placement. The GPF-DIA window scheme was also generated in EncyclopeDIA using the GPF-specific mass range and number of windows-25.

#### Spectral library

A Prosit predicted spectral library was generated for the mouse proteome (https://www.proteomicsdb.org/prosit/) ^59^. The input for the Prosit predicted spectral library was created using EncyclopeDIA’s “Convert FASTA to Prosit CSV” wizard using the complete mouse proteome (Mouse Uniprot FASTA downloaded 2019-08-21; 17015 entries). The charge range was set to 2-3, maximum missed cleavage was set to 1, m/z range: 396.4-1002.7; Default NCE-35; Default Charge-3.

#### DIA data processing

DIA raw files were converted to mzML and demultiplexed using MSConvert ^60,61^(ProteoWizard version 3.0.20084.721dd2c95) with 10 ppm accuracy and searched using EncyclopeDIA (v 0.9.0) ^56,57^ using default settings (Normal target/decoy approach, non-overlapping DIA, fragmentation set to CID/HCD (B/Y ions), 10 ppm precursor mass tolerance, 10 ppm fragment mass tolerance, 10 ppm library mass tolerance, Percolator v3-01, minimum of 3 quantitative ions).

#### Data analysis

Peptides below 1% FDR were used for the analysis. Peptide abundances were log_2_-transformed and normalized by equalizing the medians. Peptides with a coefficient of variation below 50% were used for protein estimation (see below). Protein abundance summarization was performed using Tukey’s median polish ^62^. Statistical analysis was performed using a Two-way ANOVA and a pairwise comparison using a TukeyHSD post hoc test. All p-values reported were corrected for multiple hypothesis testing. All of the analysis was performed using R (version 4.0.0).

#### Coefficient of variation

Technical replicates of the pooled sample were acquired throughout the acquisition of the experimental samples. Technical replicates are used to calculate peptide level coefficient of variation. Peptides with a CV less than 50% were used to estimate protein abundance.

#### Data availability

The DIA raw files, FASTA, Prosit library, EncyclopeDIA Chromatogram library and quant report are available at MassIVE (identifier forthcoming) and file descriptions are listed in Supplemental Table 5.

### Real-time PCR

Using all samples from the 39-week cohort, total RNA was isolated from frozen livers under Trizol (Invitrogen, CA, USA) conditions. To ensure RNA free of genomic DNA all samples underwent DNase treatment (Roche, Germany) according to the manufacture’s instructions; briefly, 1500 ng of RNA isolated from livers were treated with 1.5 μl of DNase followed by first-strand synthesis using Superscript III reverse transcriptase (Invitrogen, CA, USA) and random hexamer primers (Roche, Germany). All Real-time PCR reactions were performed using 10 uL reactions consisting of 5.0 μl of SYBR green master mix (Applied Biosystems, MA, USA), 0.2 μL of 10 μM Forward primer, 0.2μL of 10 μM Reverse primer and 4.6 μL of cDNA (Final concentration: 1.09 ng/ μL; primers listed in Supplemental Table 6). Cycle thresholds (Cts) were detected using QuantStudio 7 Flex Real-Time PCR System (Life Technologies, CA, USA). Reaction efficiency (E) was estimated for each pair of primers using a standard curve and expression levels were quantified by measuring the Cts for each sample using the E(-Cts) method. All samples were run in triplicate. Relative expression was calculated using the quantified expression from the endogenous control NONO that had stable expression levels in mouse liver across multiple samples and experimental groups.

### Western Blot

Using all liver samples from the 12- and 39-week cohorts, protein analysis was performed by Western Blot. Tissues were homogenized in RIPA extraction buffer (Cell Signaling Technology, Danvers, MA, USA) supplemented with cOmplete EDTA-free Protease inhibitor cocktail (Roche, Germany) in a cold-room using a Diagenode Bioruptor UCD-200 at HI for 15 min (30s on, 30s off) and a controlled temperature of 4°C. Tissue lysates were cleared by centrifugation at 10,000*g* for 10 min at 4°C and supernatants were transferred to new tubes. Protein concentrations were quantified by Pierce BCA protein assay kit (ThermoFisher Scientific, Waltham, MA, USA) and 20 μg of protein lysate was used per sample. 2X Laemli Sample Buffer (BioRad, Hercules, CA, USA) with 5% 2-Mercaptoethanol (BioRad, Hercules, CA, USA) were added to protein lysates and denatured at 90°C for 5 min. Samples were loaded on a NuPAGE 4-12% Bis-Tris 1.5mm Mini Protein Gel (ThermoFisher Scientific, Waltham, MA, USA) alongside with PageRuler Plus Prestained Protein Ladder 10 to 250 kDa (ThermoFisher Scientific, Waltham, MA, USA) and run using NuPAGE MOPS SDS buffer (ThermoFisher Scientific, Waltham, MA, USA). The gels were transferred onto a PVDF Immobilon-P membrane (MilliporeSigma, Burlington, MA, USA) for 90 min at 400 mA constant current. The membranes were blocked for 1h in 5% nonfat dairy milk in TBS-T (TBS Bio Rad with 0.1% Tween-20; BioRad, Hercules, CA, USA). After blocking the membranes were cut to separate the runs according to protein molecular weight of interest and each section was probed with primary antibodies diluted in 5% nonfat dairy milk in TBS-T for GAPDH (1:5,000, Cell Signalling) and either SREBP-1 (1:1,000; Abcam, Cambridge, UK), SREBP-2 (1:1,000; Abcam, Cambridge, UK) or HMGCR (1:1,000; ThermoFisher Scientific, Waltham, MA, USA) at 4°C overnight. Membranes were subsequently probed with HRP-conjugated secondary antibody (Invitrogen, CA, USA) for 2 h at room temperature. Prior to imaging, membranes were exposed to Millipore Immobilon Western HRP (MilliporeSigma, Burlington, MA, USA) substrate for 20 seconds. Bands were visualized and analyzed using an Amersham Imager 600 (GE Healthcare Life Sciences, Marlborough, MA, USA). Levels of unmodified SREBP-1, SREBP-2 and HMGCR were determined relative to GAPDH levels and compared between treatments.

### Liver Histology

After fixation in 10% phosphate-buffered formalin, a portion of the left lobe was dehydrated in ethanol and xylenes, embedded in paraffin wax and cut into 5 μm cross sections. Sections were stained with hematoxylin and eosin (H&E) and sent to the University of Pennsylvania Comparative Pathology Core for pathological analyses.

### Statistical Analyses

Statistical differences in the mean between groups were tested by two-tailed t-test for each time point and the F-test was used to test the equality of variance between groups. Overrepresentation tests were performed with PANTHER (pantherdb.org) using the GO biological processes annotation data set (released 2020-06-01) ^63,64^. The correlations of placental DNA methylation and offspring phenotypes were determined using Pearson’s correlation.

## Acknowledgements

We thank Paula Stein, Teri Ord, Monica Mainigi, and Christos Coutifaris for their technical advice with IVF procedures and helpful comments, Chris Krapp and Joanne Thorvaldsen for their technical expertise with molecular protocols. We also thank Xiaoyan Yin, Jennifer Rojas and the Rodent Metabolic Phenotyping Core (supported in part by Penn Diabetes Research Center grant (P30-DK19525)) for performing cardiometabolic assays. We acknowledge Charles-Antoine Assenmacher and the Comparative Pathology Core for performing liver pathology assessment.

## Author Contributions

LN, BAG, RMS and MSB designed the research; LN, EARC, LAV, JB and DKN performed the research; LN, EARC, LAV, JB, DKN, and YL analyzed data; LN, RMS and MSB wrote the manuscript.

---

**Supplemental Figure 1.**
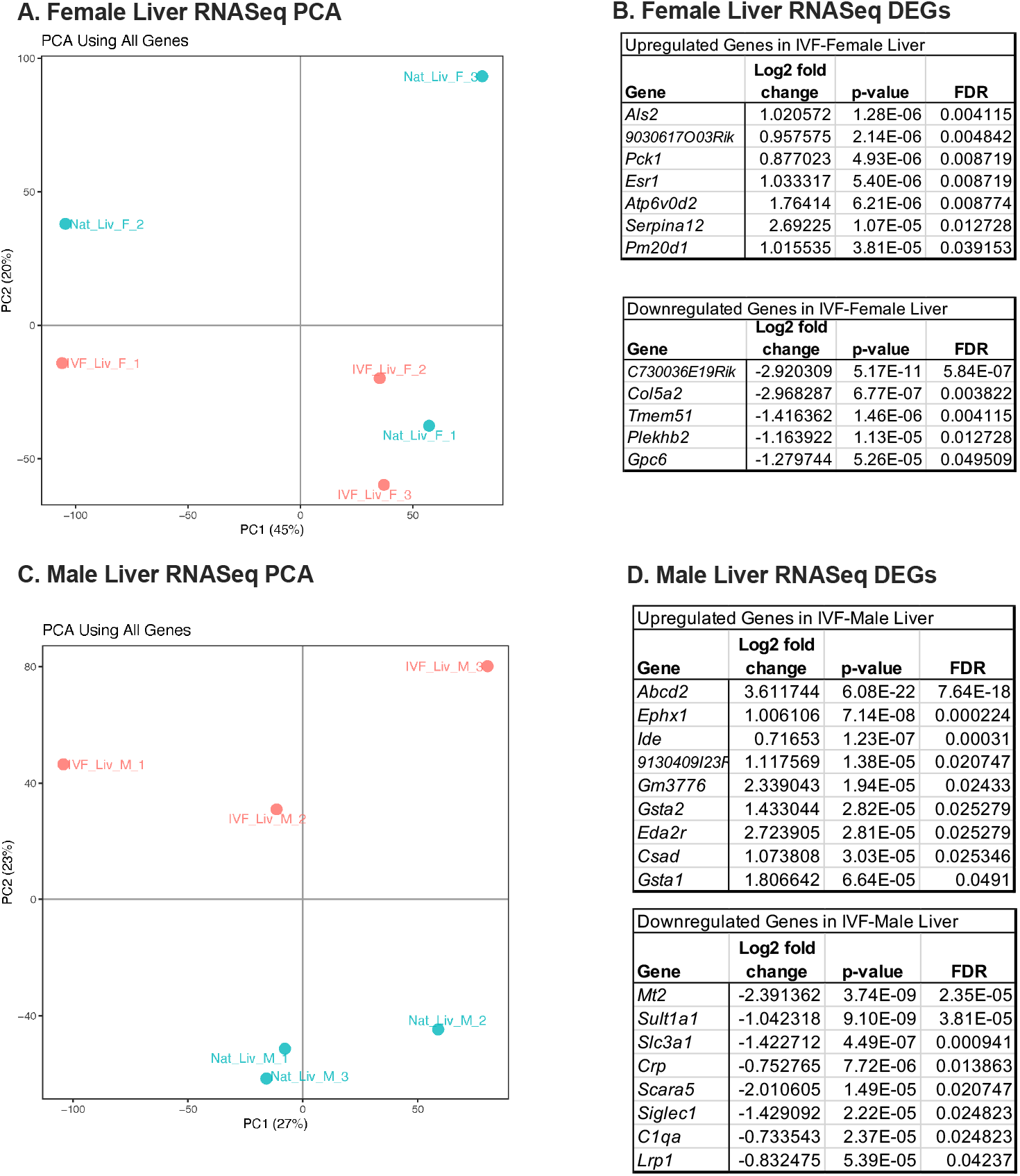
Prinicipal Component Analysis in female (**A**) and male (**C**) liver obtained through RNA-sequencing. Differentially expressed genes in female (**B**) and male (**D**) liver at 39-weeks.

**Supplemental Figure 2.**
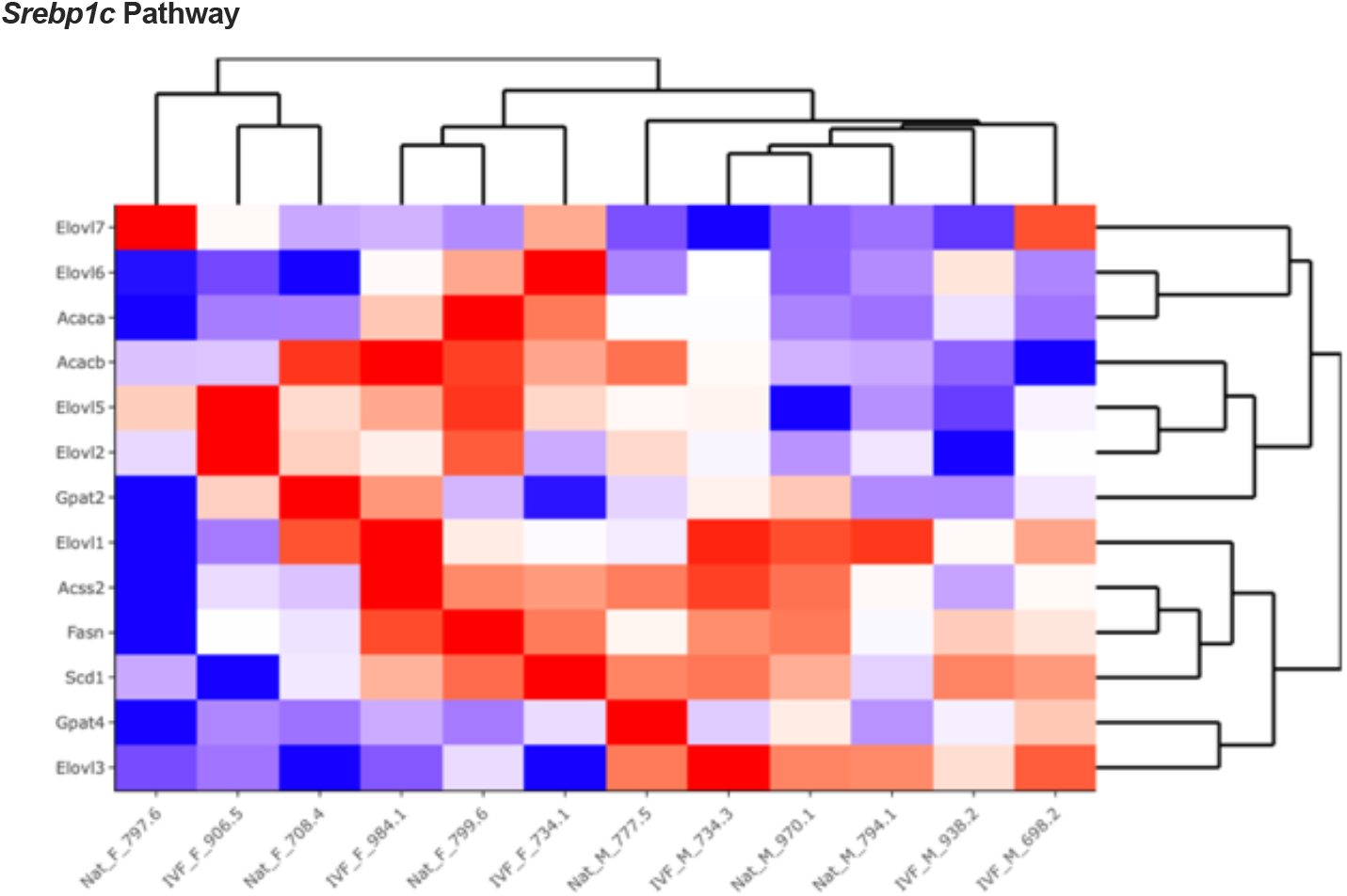
Heatmap of log_2_-transformed expression levels obtained from RNA-Seq of genes within the *Srebp-1c* pathway

**Supplemental Figure 3.**
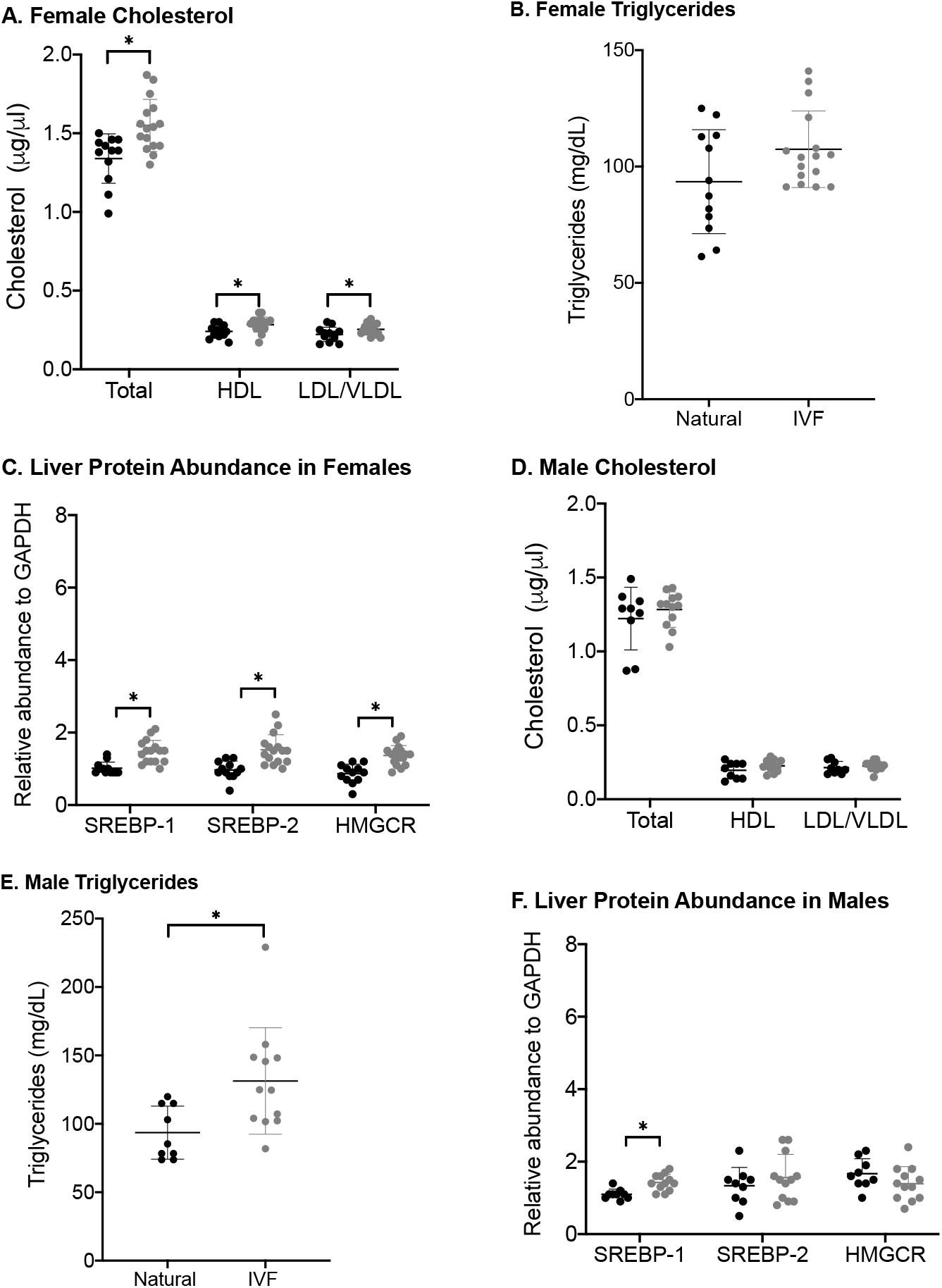
Replication of metabolic phenotype and liver protein abundance in 12-week cohort. Each data point represents an individual (n=12 Nat-female, n=15 IVF-female, n=9 Nat-male, 11-IVF-male). Total, HDL and LDL/VLDL cholesterol measured from serum in females **(A)** and males **(D)**. Serume triglycerides in females **(B)** and males **(E)**. Protein abundance of SREBP1, SREPB2 and HMGCR obtained through Western Blot in females **(C)** and males **(F)**. Gray dots indicate IVF-conceived animals. Statistical significance was determined using student’s t-test. P-value <0.05 was considered significant (*).

**Supplemental Figure 4.**
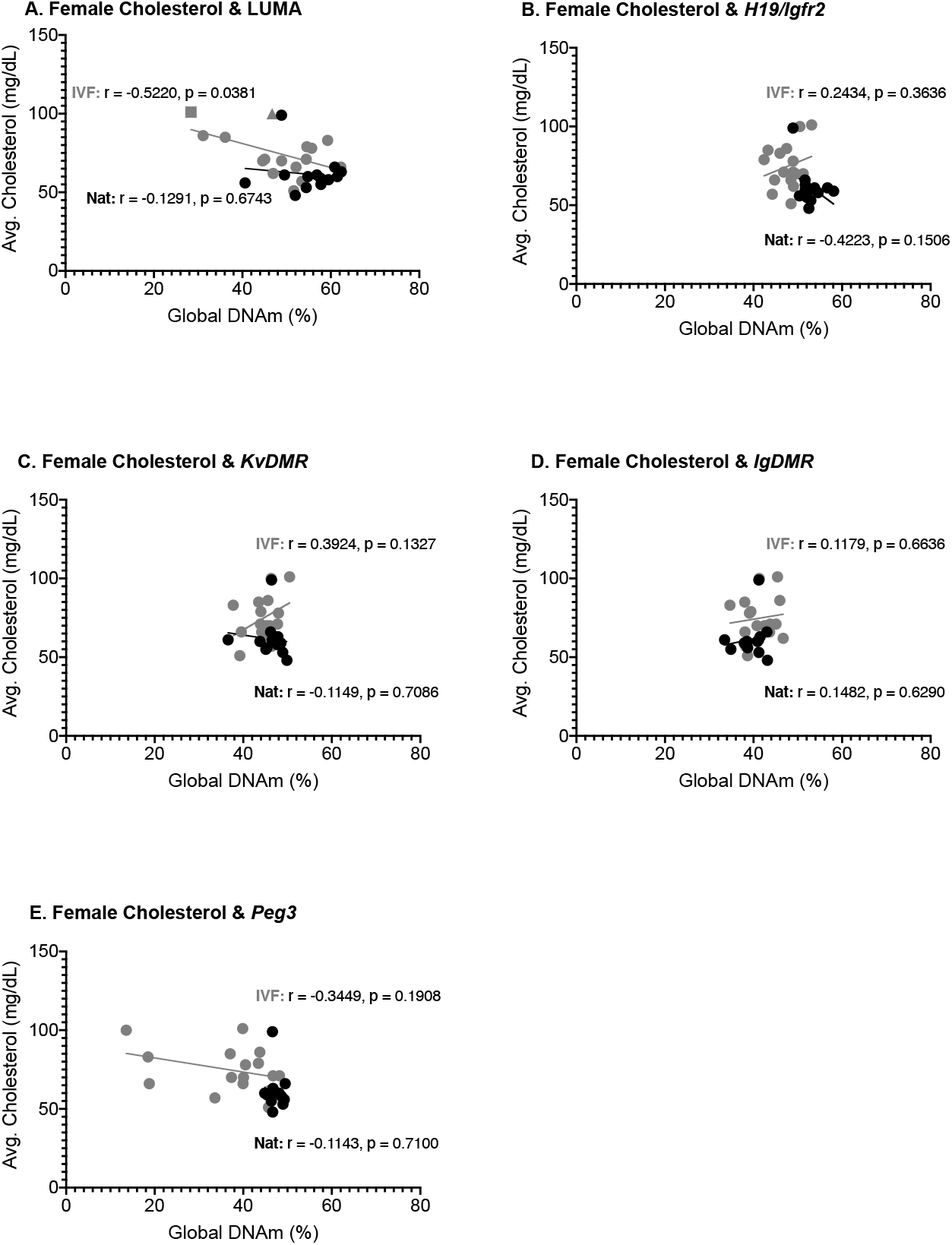
Placental DNA methylation and cholesterol. Pearson corrleation plots of average total cholesterol and global DNA methylation **(A)** and imprinting control region DNA methylation at *H19/Igf2* **(B)**, *KvDMR* **(C)**, *IgDMR* **(D)**, and *Peg3* **(E)**. Gray dots indicate IVF-conceived concepti.

**Supplemental Table 1.** Fasting serum measures from 12-39 weeks in naturally-conceived and IVF-conceived offspring.

| Supplemental Table 1 |  |  |  |  |  |  |  |  |  |  |  |  |  |  |  |  |  |  |
| --- | --- | --- | --- | --- | --- | --- | --- | --- | --- | --- | --- | --- | --- | --- | --- | --- | --- | --- |
|  | 12-weeks |  |  | 16-weeks |  |  | 20-weeks |  |  | 24-weeks |  |  | 28-weeks |  |  | 39-weeks |  |  |
|  | Nat | IVF | P | Nat | IVF | P | Nat | IVF | P | Nat | IVF | P | Nat | IVF | P | Nat | IVF | P |
| Total cholesterol (mg/dL) |  |  |  |  |  |  |  |  |  |  |  |  |  |  |  |  |  |  |
| Female |  |  |  |  |  |  |  |  |  |  |  |  |  |  |  |  |  |  |
| N | 8 | 14 |  | 13 | 15 |  | 13 | 16 |  | 12 | 16 |  | 13 | 12 |  | 11 | 15 |  |
| Mean | 66.5 | 78.2 | 0.011 | 64.4 | 82.5 | 0.006 | 56.9 | 73.6 | 0.002 | 58.9 | 71.6 | 0.009 | 65.7 | 70.1 | 0.736 | 52.93 | 71.04 | 0.033 |
| SD | 9.04 | 9.02 |  | 8.16 | 20.62 |  | 10.86 | 15.58 |  | 5.72 | 16.13 |  | 38.50 | 23.45 |  | 12.51 | 27.01 |  |
| Male |  |  |  |  |  |  |  |  |  |  |  |  |  |  |  |  |  |  |
| N | 10 | 15 |  | 12 | 18 |  | 12 | 19 |  | 12 | 19 |  | 12 | 17 |  | 12 | 17 |  |
| Mean | 106.2 | 105.9 | 0.960 | 109.1 | 100.4 | 0.133 | 99.3 | 108.9 | 0.099 | 107.4 | 104.2 | 0.634 | 108.97 | 109.33 | 0.953 | 92.4 | 123.7 | 0.015 |
| SD | 17.53 | 14.10 |  | 16.42 | 12.20 |  | 10.88 | 20.22 |  | 18.59 | 16.16 |  | 11.16 | 21.15 |  | 25.75 | 38.98 |  |
| Total triglycerides (mg/dL) |  |  |  |  |  |  |  |  |  |  |  |  |  |  |  |  |  |  |
| Female |  |  |  |  |  |  |  |  |  |  |  |  |  |  |  |  |  |  |
| N | 8 | 14 |  | 13 | 15 |  | 13 | 16 |  | 12 | 16 |  | 13 | 12 |  | 12 | 15 |  |
| Mean | 68.6 | 66.3 | 0.821 | 81.0 | 64.1 | 0.033 | 64.8 | 70.2 | 0.472 | 83.4 | 64.2 | 0.067 | 60.9 | 72.2 | 0.162 | 64.8 | 64.5 | 0.975 |
| SD | 18.3 | 28.4 |  | 22.5 | 22.53 |  | 18.9 | 20.8 |  | 28.2 | 22.8 |  | 18.4 | 20.4 |  | 24.3 | 20.6 |  |
| Male |  |  |  |  |  |  |  |  |  |  |  |  |  |  |  |  |  |  |
| N | 10 | 15 |  | 12 | 18 |  | 12 | 19 |  | 12 | 19 |  | 12 | 17 |  | 12 | 17 |  |
| Mean | 64.9 | 90.6 | 0.010 | 69.8 | 95.1 | 0.189 | 55.5 | 104.5 | 0.001 | 78.0 | 90.3 | 0.345 | 67.2 | 97.7 | 0.029 | 75.2 | 96.3 | 0.142 |
| SD | 31.3 | 12.7 |  | 17.3 | 37.2 |  | 15.0 | 54.0 |  | 31.0 | 40.0 |  | 19.1 | 49.0 |  | 29.3 | 45.6 |  |
| Insulin (ng/mL) |  |  |  |  |  |  |  |  |  |  |  |  |  |  |  |  |  |  |
| Female |  |  |  |  |  |  |  |  |  |  |  |  |  |  |  |  |  |  |
| N | 7 | 13 |  | 12 | 13 |  | 13 | 16 |  | 12 | 16 |  | 12 | 12 |  | 11 | 14 |  |
| Mean | 0.12 | 0.13 | 0.847 | 0.18 | 0.16 | 0.819 | 0.16 | 0.13 | 0.728 | 0.12 | 0.16 | 0.324 | 0.09 | 0.16 | 0.061 | 0.22 | 0.42 | 0.044 |
| SD | 0.13 | 0.08 |  | 0.13 | 0.18 |  | 0.26 | 0.11 |  | 0.08 | 0.13 |  | 0.08 | 0.08 |  | 0.22 | 0.24 |  |
| Male |  |  |  |  |  |  |  |  |  |  |  |  |  |  |  |  |  |  |
| N | 9 | 13 |  | 12 | 17 |  | 12 | 19 |  | 11 | 19 |  | 12 | 17 |  | 12 | 17 |  |
| Mean | 0.32 | 0.22 | 0.493 | 0.22 | 0.39 | 0.069 | 0.18 | 0.39 | 0.047 | 0.13 | 0.58 | <0.001 | 0.23 | 0.55 | <0.001 | 0.50 | 1.09 | 0.012 |
| SD | 0.40 | 0.14 |  | 0.13 | 0.34 |  | 0.17 | 0.37 |  | 0.05 | 0.39 |  | 0.12 | 0.30 |  | 0.28 | 0.83 |  |
| Glucose (mg/dL) |  |  |  |  |  |  |  |  |  |  |  |  |  |  |  |  |  |  |
| Female |  |  |  |  |  |  |  |  |  |  |  |  |  |  |  |  |  |  |
| N | 8 | 14 |  | 13 | 15 |  | 13 | 16 |  | 13 | 16 |  | 13 | 12 |  | 12 | 15 |  |
| Mean | 91.6 | 88.4 | 0.391 | 89.1 | 83.7 | 0.269 | 97.2 | 87.0 | 0.038 | 93.4 | 87.3 | 0.608 | 86.9 | 91.9 | 0.388 | 97.0 | 94.5 | 0.579 |
| SD | 11.7 | 13.9 |  | 15.2 | 8.5 |  | 13.6 | 10.6 |  | 13.2 | 14.1 |  | 11.6 | 16.5 |  | 12.7 | 9.3 |  |
| Male |  |  |  |  |  |  |  |  |  |  |  |  |  |  |  |  |  |  |
| N | 10 | 15 |  | 12 | 18 |  | 12 | 19 |  | 12 | 19 |  | 12 | 17 |  | 12 | 17 |  |
| Mean | 82.5 | 93.3 | 0.061 | 87.0 | 102.5 | 0.001 | 97.7 | 109.8 | 0.075 | 95.9 | 117.4 | <0.001 | 108.0 | 109.7 | 0.781 | 123.3 | 130.7 | 0.391 |
| SD | 10.5 | 16.8 |  | 10.3 | 13.7 |  | 16.57 | 19.7 |  | 14.4 | 14.5 |  | 18.1 | 10.7 |  | 22.1 | 22.8 |  |

**Supplemental Table 2.**
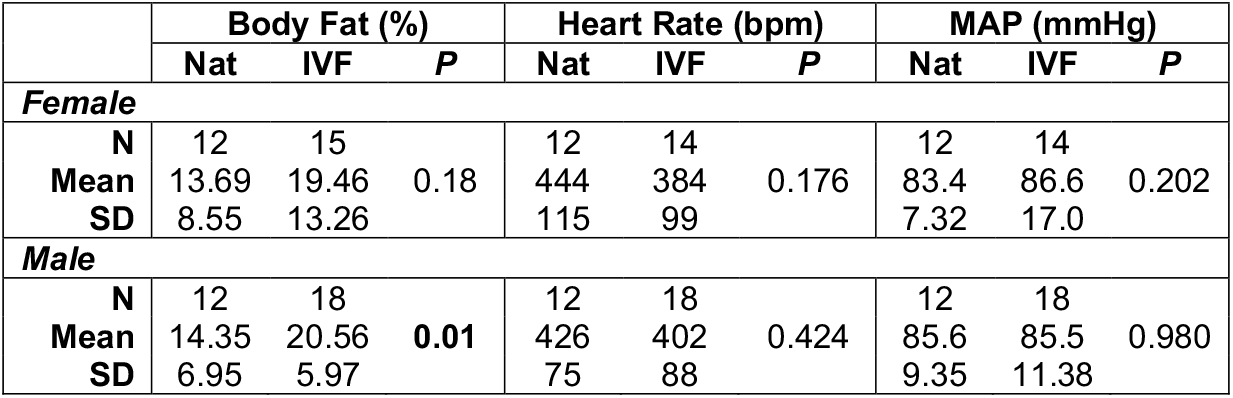
Percent body fat, heart rate and mean arterial pressure (MAP) in naturally-conceived and IVF-conceived offspring at 39-weeks-of-aαe.

**Supplemental Table 3.** RNA-seq reads, alignment and mapping in 39-week liver samples.

| **Supplemental Table 3** |  |  |  |  |  |  |  |
| --- | --- | --- | --- | --- | --- | --- | --- |
| *RNA-seq reads, alignment and mapping in 39-week liver samples.* |  |  |  |  |  |  |  |
| **Sample** | **Raw read pairs** | **After trimming** | **Uniquely mapped** | **Mapped to multiple loci** | **Total mapped** | **Mapping rate** | **Total fragments assigned to RefSeq gene** |
| IVF_Liv_F_1 | 17,426,970 | 15,042,744 | 9,659,881 | 2,838,754 | 12,498,635 | 83% | 6,826,534 |
| IVF_Liv_F_2 | 33,869,890 | 32,426,033 | 24,045,557 | 2,881,262 | 26,926,819 | 83% | 21,833,725 |
| IVF_Liv_F_3 | 31,766,777 | 30,656,420 | 23,550,760 | 3,014,176 | 26,564,936 | 87% | 22,073,863 |
| IVF_Liv_M_1 | 20,605,095 | 19,964,051 | 14,601,536 | 2,492,574 | 17,094,110 | 86% | 14,002,847 |
| IVF_Liv_M_2 | 32,775,511 | 31,781,775 | 21,679,285 | 4,838,267 | 26,517,552 | 83% | 21,046,484 |
| IVF_Liv_M_3 | 33,088,796 | 31,894,643 | 21,507,026 | 5,486,532 | 26,993,558 | 85% | 21,762,008 |
| Nat_Liv_F_1 | 33,938,663 | 32,912,162 | 24,778,235 | 3,359,634 | 28,137,869 | 85% | 23,430,136 |
| Nat_Liv_F_2 | 26,189,256 | 22,088,064 | 13,446,688 | 4,686,204 | 18,132,892 | 82% | 8,056,230 |
| Nat_Liv_F_3 | 26,810,170 | 25,870,427 | 16,310,747 | 2,791,363 | 19,102,110 | 74% | 14,302,287 |
| Nat_Liv_M_1 | 36,452,150 | 35,178,250 | 24,398,639 | 4,615,618 | 29,014,257 | 82% | 24,235,375 |
| Nat_Liv_M_2 | 34,686,518 | 33,485,511 | 22,521,094 | 5,484,424 | 28,005,518 | 84% | 23,448,074 |
| Nat_Liv_M_3 | 39,669,372 | 38,253,629 | 27,176,336 | 5,218,226 | 32,394,562 | 85% | 27,832,271 |

**Supplemental Table 4.** Description of mass spectrometry raw files

| **FileName** | **BioRep** | **TechRep** | **Method** |
| --- | --- | --- | --- |
| 20200311_HFX_Bartolomei_IVF-Liver_Female-IVF-01_16.raw | 1 | 1 | DIA |
| 20200311_HFX_Bartolomei_IVF-Liver_Female-IVF-02_22.raw | 2 | 1 | DIA |
| 20200311_HFX_Bartolomei_IVF-Liver_Female-IVF-03_36.raw | 3 | 1 | DIA |
| 20200311_HFX_Bartolomei_IVF-Liver_Female-IVF-04_38.raw | 4 | 1 | DIA |
| 20200311_HFX_Bartolomei_IVF-Liver_Female-IVF-05_43.raw | 5 | 1 | DIA |
| 20200311_HFX_Bartolomei_IVF-Liver_Female-Nat-06_18.raw | 6 | 1 | DIA |
| 20200311_HFX_Bartolomei_IVF-Liver_Female-Nat-07_21.raw | 7 | 1 | DIA |
| 20200311_HFX_Bartolomei_IVF-Liver_Female-Nat-08_35.raw | 8 | 1 | DIA |
| 20200311_HFX_Bartolomei_IVF-Liver_Female-Nat-09_37.raw | 9 | 1 | DIA |
| 20200311_HFX_Bartolomei_IVF-Liver_Female-Nat-10_45.raw | 10 | 1 | DIA |
| 20200311_HFX_Bartolomei_IVF-Liver_Male-IVF-11_15.raw | 11 | 1 | DIA |
| 20200311_HFX_Bartolomei_IVF-Liver_Male-IVF-12_20.raw | 12 | 1 | DIA |
| 20200311_HFX_Bartolomei_IVF-Liver_Male-IVF-13_33.raw | 13 | 1 | DIA |
| 20200311_HFX_Bartolomei_IVF-Liver_Male-IVF-14_40.raw | 14 | 1 | DIA |
| 20200311_HFX_Bartolomei_IVF-Liver_Male-IVF-15_44.raw | 15 | 1 | DIA |
| 20200311_HFX_Bartolomei_IVF-Liver_Male-Nat-16_17.raw | 16 | 1 | DIA |
| 20200311_HFX_Bartolomei_IVF-Liver_Male-Nat-17_23.raw | 17 | 1 | DIA |
| 20200311_HFX_Bartolomei_IVF-Liver_Male-Nat-18_34.raw | 18 | 1 | DIA |
| 20200311_HFX_Bartolomei_IVF-Liver_Male-Nat-19_39.raw | 19 | 1 | DIA |
| 20200311_HFX_Bartolomei_IVF-Liver_Male-Nat-20_42.raw | 20 | 1 | DIA |
| 20200311_HFX_Bartolomei_IVF-Liver_pool-pool-01_19.raw |  | 1 | DIA |
| 20200311_HFX_Bartolomei_IVF-Liver_pool-pool-02_32.raw |  | 2 | DIA |
| 20200311_HFX_Bartolomei_IVF-Liver_pool-pool-03_41.raw |  | 3 | DIA |
| 20200311_HFX_Bartolomei_IVF-Liver_pool-pool-04_46.raw |  | 4 | DIA |
| 20200311_HFX_Bartolomei_IVF-Liver_GPF-01_25.raw |  |  | GPF |
| 20200311_HFX_Bartolomei_IVF-Liver_GPF-02_26.raw |  |  | GPF |
| 20200311_HFX_Bartolomei_IVF-Liver_GPF-03_27.raw |  |  | GPF |
| 20200311_HFX_Bartolomei_IVF-Liver_GPF-04_28.raw |  |  | GPF |
| 20200311_HFX_Bartolomei_IVF-Liver_GPF-05_29.raw |  |  | GPF |
| 20200311_HFX_Bartolomei_IVF-Liver_GPF-06_30.raw |  |  | GPF |

**Supplemental Table 5.**
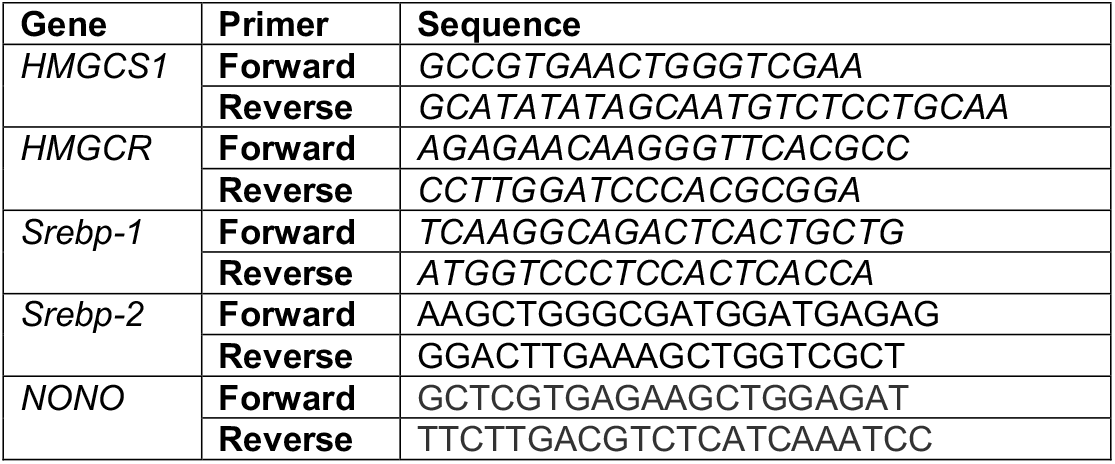
Real-time PCR primers

**Supplemental Table 6.** Percent DNAm in naturally-conceived and IVF-conceived term placenta.

| Supplemental Table 6 |  |  |  |  |  |  |  |  |  |  |  |  |  |  |  |
| --- | --- | --- | --- | --- | --- | --- | --- | --- | --- | --- | --- | --- | --- | --- | --- |
| Percent DNAm in naturally-conceived and IVF-conceived term placenta. |  |  |  |  |  |  |  |  |  |  |  |  |  |  |  |
|  | Global DNAm |  |  | H19/Igf2 |  |  | KvDMR1 |  |  | IgfDMR |  |  | Peg3 |  |  |
|  | Nat | IVF | P | Nat | IVF | P | Nat | IVF | P | Nat | IVF | P | Nat | IVF | P |
| Female |  |  |  |  |  |  |  |  |  |  |  |  |  |  |  |
| N | 13 | 14 |  | 13 | 16 |  | 13 | 16 |  | 13 | 16 |  | 13 | 16 |  |
| Mean | 55.13 | 48.20 | 0.026 | 52.97 | 47.70 | <0.001 | 46.14 | 44.51 | 0.202 | 39.54 | 41.33 | 0.146 | 47.30 | 37.08 | <0.001 |
| SD | 6.15 | 9.55 |  | 2.47 | 2.96 |  | 3.32 | 3.35 |  | 2.93 | 3.51 |  | 1.47 | 10.77 |  |
| Male |  |  |  |  |  |  |  |  |  |  |  |  |  |  |  |
| N | 12 | 17 |  | 12 | 18 |  | 12 | 19 |  | 12 | 18 |  | 12 | 18 |  |
| Mean | 52.75 | 53.45 | 0.803 | 52.59 | 52.05 | 0.724 | 46.77 | 45.15 | 0.429 | 38.36 | 41.05 | 0.079 | 38.36 | 41.05 | 0.002 |
| SD | 6.26 | 8.77 |  | 4.27 | 3.62 |  | 5.49 | 5.41 |  | 18.59 | 5.0 |  | 3.08 | 6.72 |  |

